# Modelling time evolution of in blood alcohol concentration

**DOI:** 10.1101/2021.07.25.452934

**Authors:** Antoni Oliver Gelabert

## Abstract

A theoretical model based on mass conservation is proposed to predict blood alcohol concentration (BAC) over time. The proposed model takes as input the body parameters, the data timing for each drink, as well as his alcoholic strength and ingested amount. Three test cases with variations are presented and discussed. The results given are obtained by solving numerically the equation proposed. The solution predicts a well located alcohol peak and a reasonable time of alcohol body cleaning. This model represents a first order approach to a complex problem. Similar scenarios, like, for instance, drug concentrations can be modelled by using the same concept. The proposed model has been implemented on easy-to-use program COGORZA, which source code is available online at https://github.com/tonibois/Cogorza.

## 1 Introduction

Early studies have been conducted for the purpose of predicting BAC [1], [2]. Recently, a machine learning was proposed and trained using user data from a smart breathalyzer [3]. However, there is still a lack of connectivity between ingestion rates that determine the evolution of the curve with the resulting BAC itself using simple and intelligible model.

In this work I present a mathematical formulation to model and predict an approximation of the concentration evolution of a substance in the bloodstream. The proposed concept is based on a single equation with arbitrary input drinks modelled as a set of step-wise functions that equally distributes injected alcohol mass of the drink over the drinking time. Timing in terms of consumption is a key, commonly forgotten due to its modeling complexity. An accurate prediction model must have at least the information of the time of intake (measured from the time elapsed since the first intake), the type of drink (especially its strength) and the amount of alcohol ingested and absorbed into the bloodstream. Then, a drink that is consumed faster will produce higher input value with shorter time than a drink that is consumed slowly.

In this model, there’s assumed a 10% and 12% of alcohol absorption for man and woman, respectively, which is take from literature [1] and is further explained below. The rate of absorption depends on the amount and the kind of previous food ingested by the subject, but is between 10 and 60 min^2^. A constant rate of alcohol expulsion is set to 12 g/L^3^. The mass of the drinker are also included in the formulation of the model, so it relates with total blood amount. The simulated scenarios are presented as hypothetical situations in which the taking decision phase becomes critical. The results provides direct information about approximate time and magnitude of the BAC peak and the required time to cleaning the body out of alcohol.

In order to aim the reader to check and experiment with that proposal, a program has been developed in both *fortran* and *python* languages. This implementations provide the graphical output and predicted time series of BAC.

## 2 Theoretical model

The model proposed in this work is based on the arbitrary alcohol intake and constant elimination rates. Firstly, alcohol mass is computed using drink volume and strength derived from drink types (for instance, shot of tequila, half beer, a cup of wine or a glass of *whiskey on the rocks*). These parameters are used to characterize the input function. Using equation 1, the mass of alcohol in a drink *m*_*i*_ is determined as a function of the strength of the drink (G), its volume (V) and density (D), which in the case of alcohol is 0.789 g/mL.

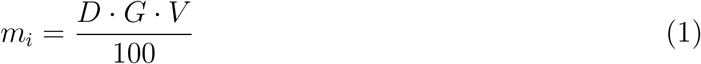

Thus a pint (V=473.2 ml) of beer (G=5%) is translated to about 19 g of alcohol. Table 1 shows the amount of alcohol associated with these parameters for different kind of drinks.

**Table 1.**
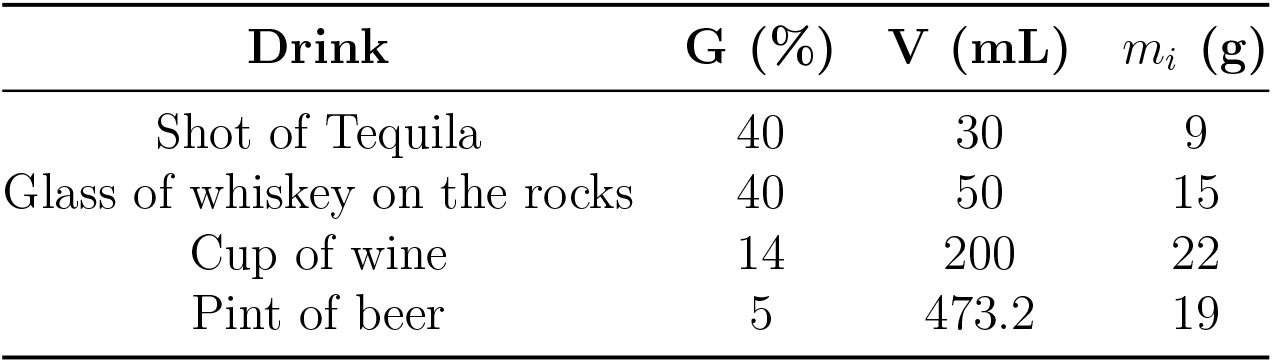
Alchol mass (*m*_*i*_) for some drinks

Once the ingested mass is obtained, the concentration that passes into the subject’s blood-stream must be calculated. To calculate the blood alcohol concentration, the amount of substance present in the blood will be determined between the amount of the subject’s blood, which is proportional to its mass, M. The value of *C*_*s*_ has been estimated in different studies [4–6] and some of those studies involves gender class, age, height and body complexity. In order to reduce the number of parameters in this study an averaged value only proportional to the mass amount is taken (*C*_*s*_ = 0.067*L/kg*). This value means that a person with 100 kg weight will have 6.7 liters of blood.

On the other hand, the proportion of alcohol that passes into the circulatory system is approximately *A*_*bs*_ *≈* 0.15 = 15%^4^,^5^. Taking these factors into account, the blood alcohol for a drink *i* (*a*_*i*_) is calculated using equation 2.

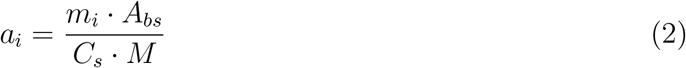

Equation 2 can be related to the parameters found in early studies [1], in which BAC is computed by equation 3.

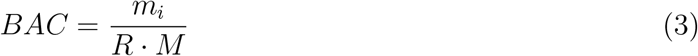

Where R is the body diffusion coefficient, which is equal to 0.55 for women and 0.68 for men [7]. Assuming that the concentration of blood per unit mass (*C*_*s*_) is the same in men and women, from the equations 2 and 3 an alcohol absorption coefficient, *A*_*bs*_ is deduced to be around 12 % for women and 10 % for men.

Taking these relationships into account, we can generate a model of alcohol intake based on step functions, Θ_*i*_(*t*) whose area under the curve is the result of the mass of alcohol ingested *m*_*i*_. The alcohol intake, therefore, will be an input function resulting from the sum of all the intakes *i*. Each intake will start at an initial time *t*_*oi*_ and end at a final time *t*_*fi*_. The time increment *δt*_*i*_ = *t*_*fi*_ *− t*_*oi*_ is the time taken to consume the ith drink.

The alcohol expends between *t*_*abs*_ = 15 to 60 minutes since is ingested to arrive at bloodstream depending of food intake degree *fid* (which range from 1, empty stomach to 4, full stomach). Therefore, the alcohol absorbed in individual drinks can be expressed as 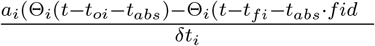 Thus, the total alcohol intake in time, *i*(*t*), due to N drinks, and assuming an initial blood alcohol concentration of *a*_*o*_ can be written as shown in equation 4.

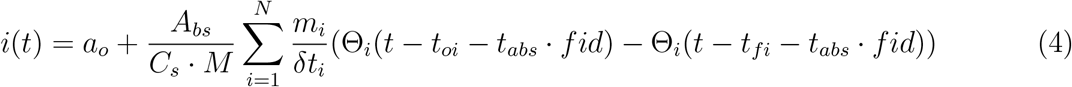

To estimate the rate of alcohol elimination in the body, a constant rate is assumed whenever there is a concentration of in blood alcohol greater than zero. That is, when alcohol is present in the body, it is eliminated at a constant rate of *e ≈* 0.12 grams of alcohol per liter of blood every hour. Using *Cs*, this is translated into a mass removal rate of alcohol in grams per second, given by *El ≈* 3.33 *·* 10^*−*5^*g/*(*L · s*).

By application of the equation of mass conservation, in blood alcohol concentration can be described as the mass that enters the body minus the mass that leaves from it using equation 5.

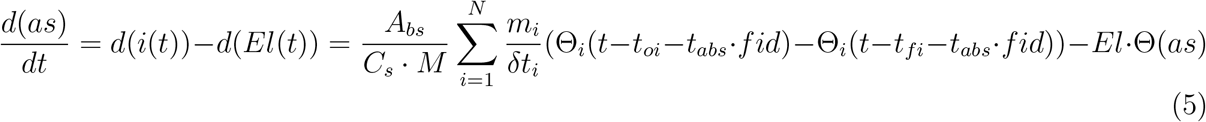

Where Θ(*as*) forces the condition that the elimination rate *El*(*t*) will be equal to zero when in blood alcohol concentration become null. By Integrating the expression 5, we obtain equation 6.

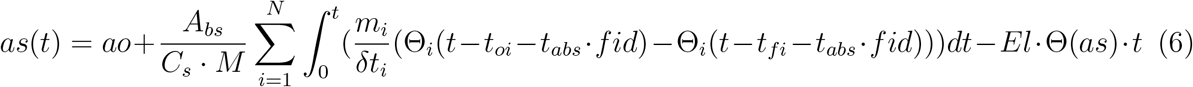

Very related to in blood alcohol concentration, as(t), is the level of alcohol in expired air (ae) that can be obtained using expression found in [8]. From this work is derived the equation 7.

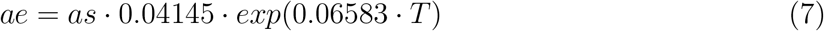

Where T is the body temperature expressed in Celsius (i.e. about 37.5°*C* under normal conditions). Then, the above equation is reduced to 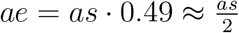. That is, the expired air alcohol level (ae) is approximately half of the blood alcohol level (as) if ae is expressed in mg/L and as in g/L. In table 2 indicative risk zones based on the degree of alcohol in blood and expired air are shown. This regions are also shown in the output given by COGORZA.

**Table 2.** Danger zones based on the level of alcohol in blood (as) and expired air (ae)

| zone | as (g/L) | ae (mg/L) |
| --- | --- | --- |
| Non-Risk | 0.0-0.3 | 0.0-0.15 |
| Happy zone | 0.3-0.8 | 0.15-0.40 |
| The dot | 0.8-1.5 | 0.40-0.75 |
| drunkenness | 1.5-3.0 | 0.75-1.5 |
| Death zone | > 3.0 | >1.5 |

## 3 Methods

Numerical solutions have been found solving equation 5 with step size of *dt* = 1 second. Simulations performed are ranging from 360 (6 h) to 1440 min (24 h) length. A male subject of 92 kg is used as the drinker profile. Time absorption is set to 20 minutes with 10% rate of absorption. The solution found for equation 6 has been smoothed by applying a 1D Gaussian filter with *σ* = 100. Further details can be found at the end of this work, where source code is made available.

## 4 Results

Following, a set of hypothetical test cases are exposed and discussed. The first case has a moderate context, that takes place in Spain. In Spain and great part of Europe, traffic law allows to fine standard drivers which BAC in alcoholic test is higher than 0.5 g/L. A male subject with M=92 kg of body mass is going out for a dinner. He consumes three half beers and one *whiskey on the rocks* during a full night.

Let us assume that the subject at 8:00 pm begins to drink a beer in *snack/vermouth* mode. He finishes the first drink after 30 minutes. Then he pauses and at the start of dinner at 9:00 p.m., when he begins drinking the second beer, which ends after 20 minutes. At 9:30 p.m. the subject starts the next drink, finishing it at 10 p.m. along with end of dinner. Once reached after-dinner stage, our subject consumes a glass of *whiskey on the rocks*. He starts that last drink at 10:20 p.m. which ends at 10:40 p.m. At 11 pm, the drinker decides to drive to home with his car. Is he prepared to drive without risk? Should him leave later? Or would it became better for him to leave earlier? The simulation that follows will use this input data to shed some light on these complex-looking questions.

By feeding this data in to the COGORZA program, we obtain the evolution curve shown in Figure 1. The solution found through COGORZA invites us to be cautious, because if we do not want our driver to be on risk, he must stay in the blue zone. As can be seen in Figure 1, immediately after the third drink, the subject passes the safe zone and enters the happy zone, where he can give a positive breath test. Therefore, if he decides to finish last whiskey, he would wait until at least 300 minutes after the first drink start time. That is, instead of leaving at 11 p.m. (i.e. at 180 minutes from first drink start time, which is very close to the BAC peak), he should stay for at least two more hours. Then, he should leave after 1 a.m. without having consumed any other drink during the last 140 minutes.

**Figure 1.**
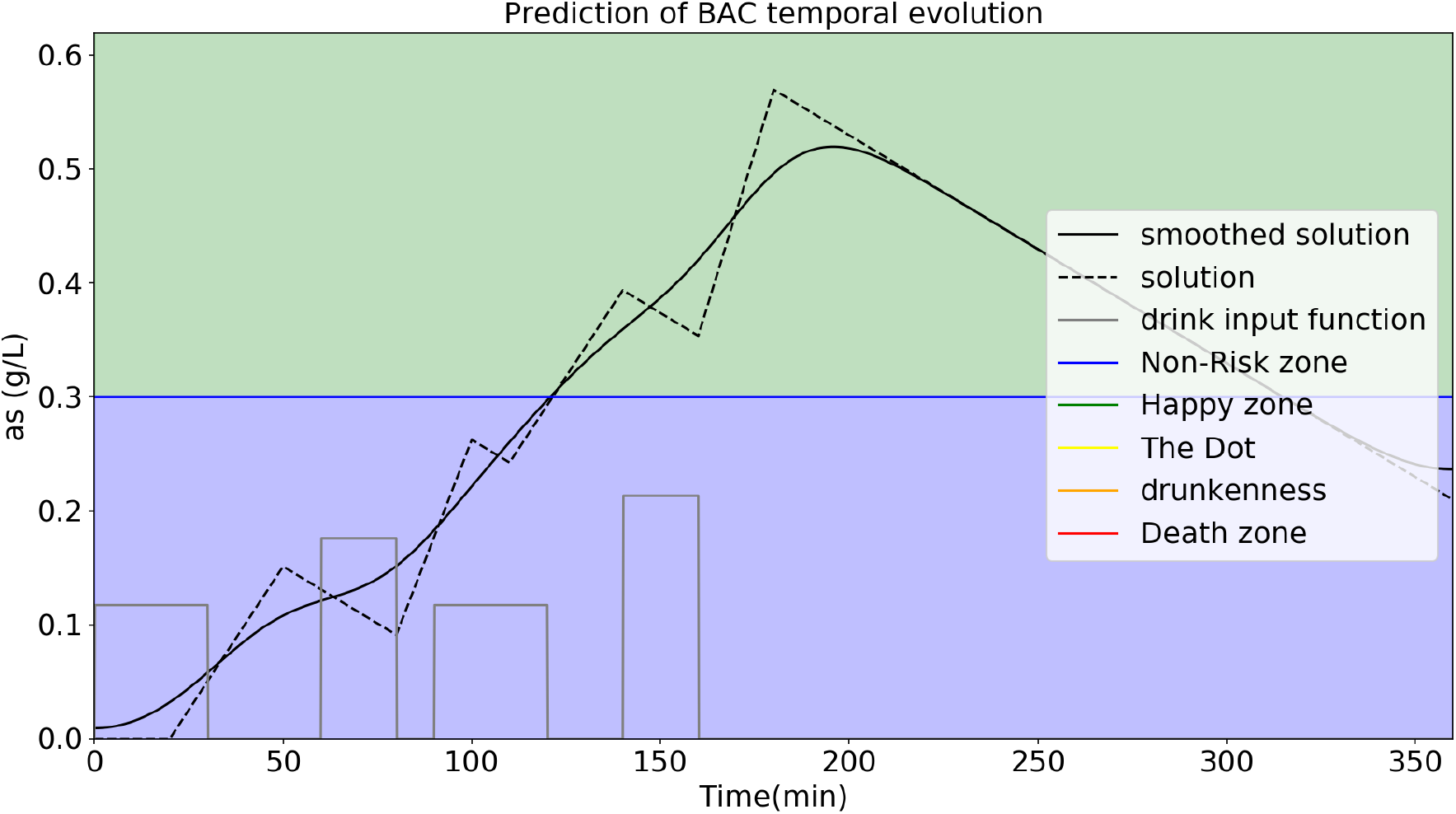
Predicted temporal evolution of BAC of moderate case number one (black dotted line). The solid black line represents the smoothed solution. The step functions in gray in the lower part represents the input drinks made. The colored areas in the background represent areas of risk due to BAC concentration levels.

The second case presented is the case of a subject with the same characteristics as the previous one, who goes out to party and takes the following shots:

1. At 4pm, he make the first drink. It’s a *cubalibre* with 60 mL of rum with a 40% graduation. At 4:25 p.m. he has finished that drink and waits for 5 minutes to start the next one.
2. At 4.30 pm the second drink begins, with identical conditions to the previous one.
3. It is 5:00 p.m. and the subject begins his third drink.
4. At 5.30 pm the fourth drink begins, which is consumed faster than the others, taking only 10 minutes for the subject for finishing it.
5. The fifth drink, like the others, starts at 5:45 p.m. and ends in only 5 minutes.
6. The sixth drink is made a little later, starting at 6.30 pm and ending at 6.55 pm.

Given this scenario, it is worth considering whether the drinker will have made an exaggerated intake of alcohol, surpassing the so-called dot area to enter the drunkenness area. Place your bets and see the solution obtained by the COGORZA program, the result of which is shown in Figure 2.

**Figure 2.**
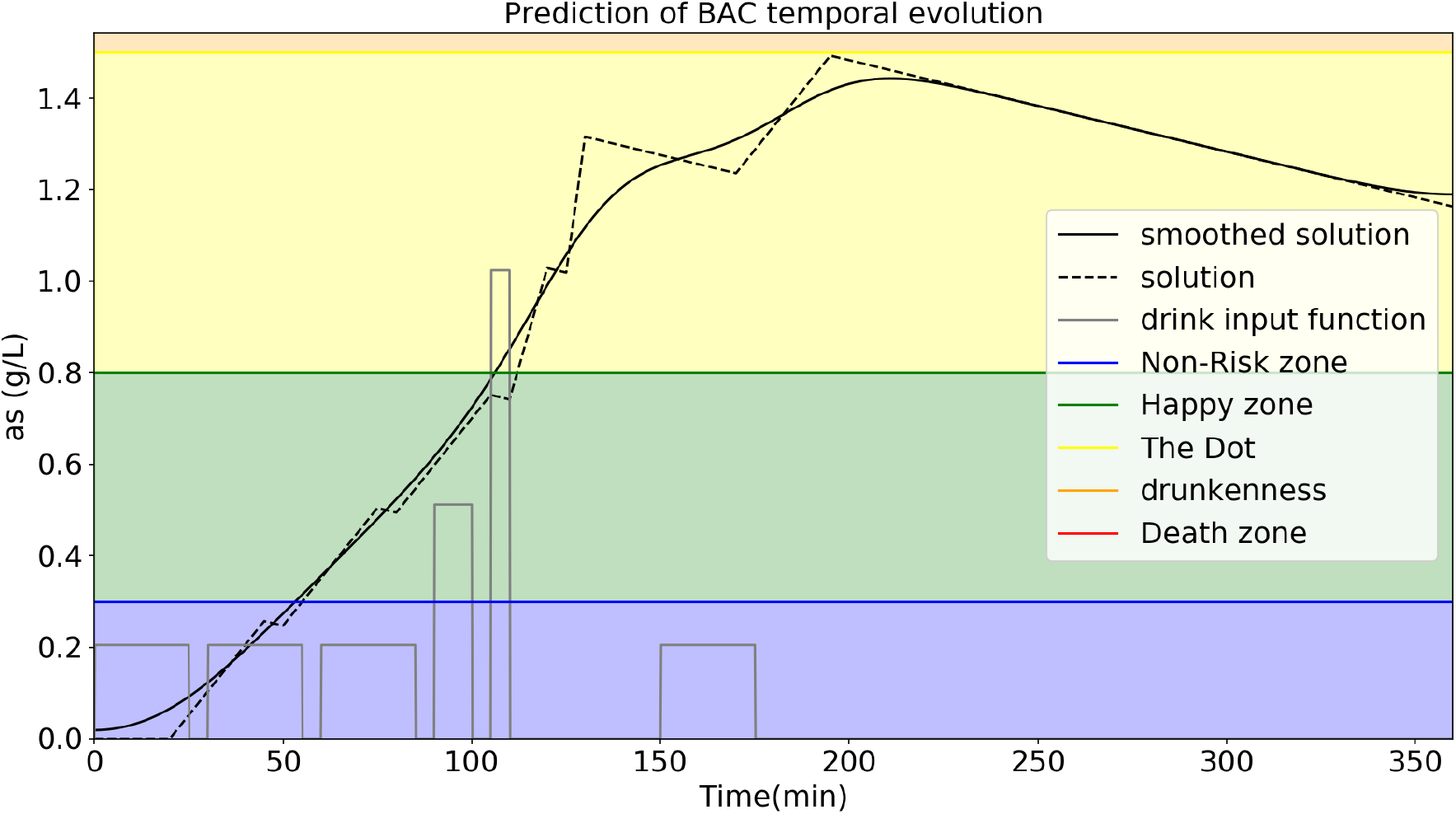
Predicted temporal evolution of BAC of moderate case number two (black dotted line). The solid black line represents the smoothed solution. The step functions in gray in the lower part represents the input drinks made. The colored areas in the background represent areas of risk due to BAC concentration levels. In that case, the body mass of the subject is high (M=92 kg).

We see that by very little, the drunkenness area (1.5 to 3.0 g/l) is not reached. However, a somewhat more critical result is obtained from the same experiment performed with a 70 kg subject (see Figure 3).

**Figure 3.**
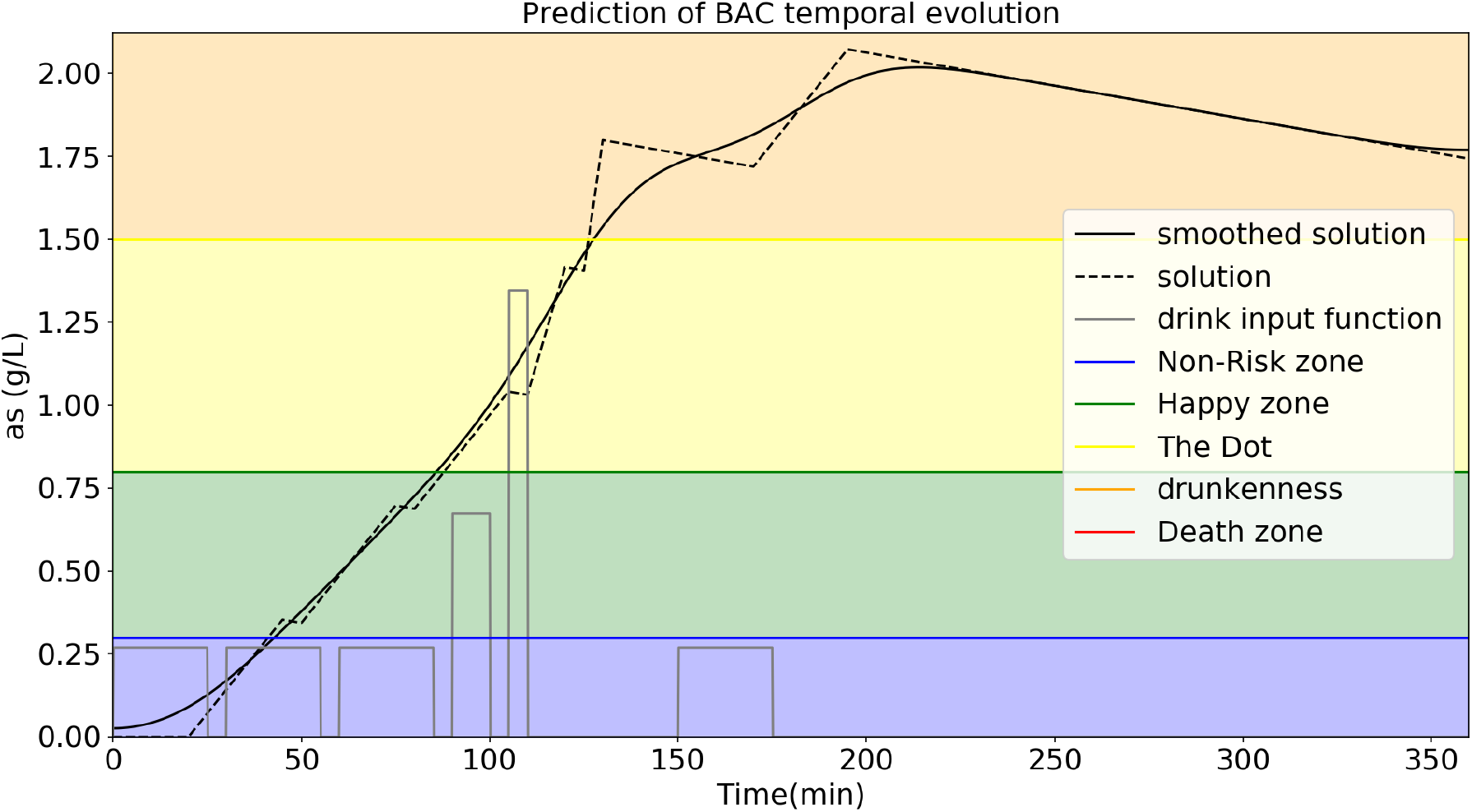
Predicted temporal evolution of BAC for moderate case number two (black dotted line). The solid black line is the smoothed solution. The step functions in gray in the lower part represents the input drinks made. Colored areas in the background represent areas of risk due to BAC concentration levels. In that case, the body mass of the subject is high (M=70 kg).

We see that the results obtained in Figure 3 indicates that this difference in weight is enough to enter the drunkenness, once the blood alcohol level of 1.5 g/l is exceeded.

In the last case of study, an identical subject to the one shown in the first two cases is used. This drinker attends at a long event and, despite the fact that the total consumption of alcohol is high, the timing of the shots determines the severity of BAC peak. The subject in question drinks twelve glasses of wine. Every hour he consumes three cups of wine and he drink without interruption, so he reaches twelve glasses of wine in 4 hours. The result of these tests is a blood alcohol peak that enters the risk zone (Figure 4).

**Figure 4.**
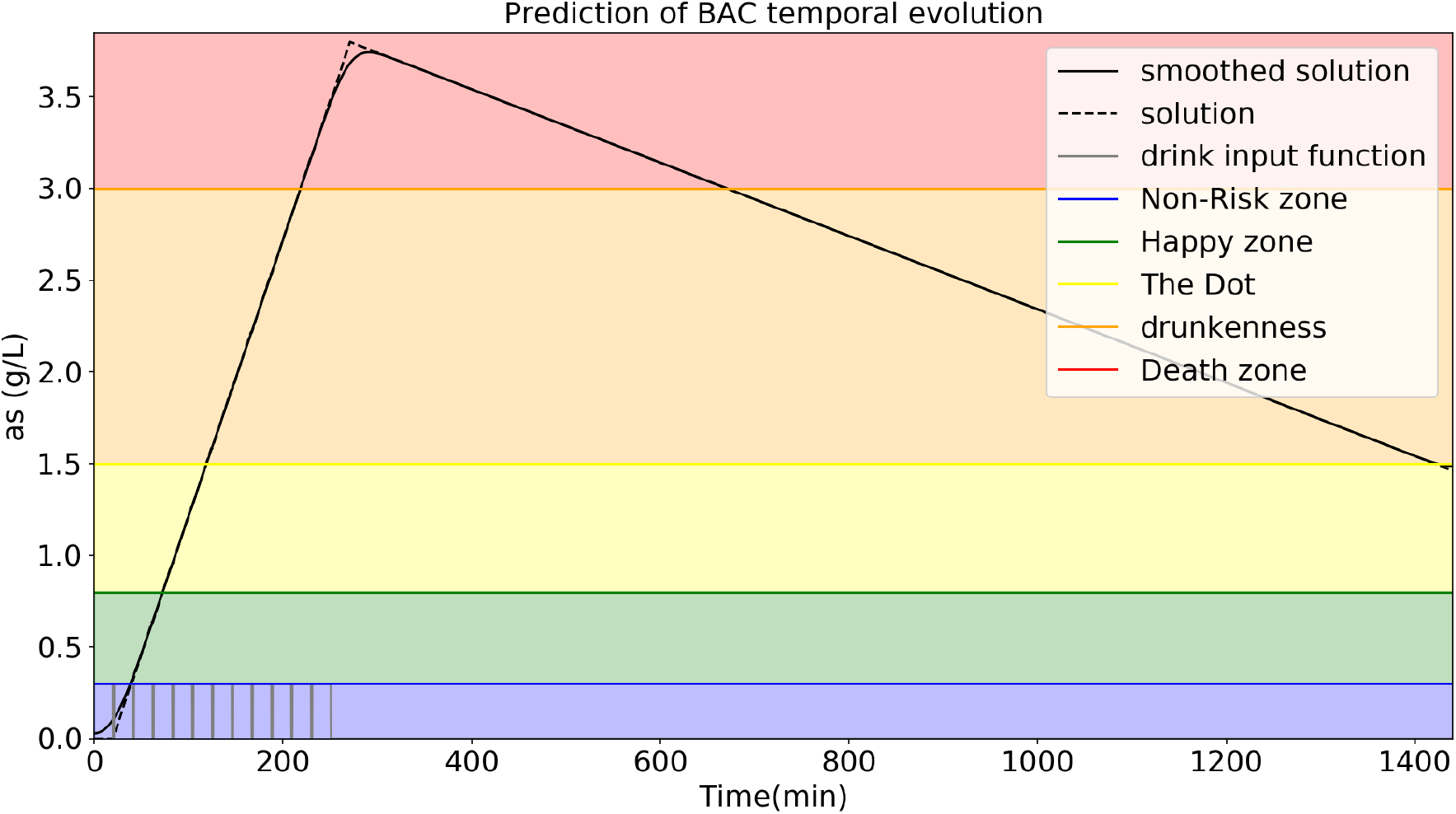
Predicted temporal evolution of BAC of the case of study number three for a drinker that drink one cup of wine every 20 minutes without interruption (black dotted line). The solid black line represents the smoothed solution. The step functions in gray in the lower part represents the input drinks made. Colored areas in the background represent areas of risk due to high BAC levels.

If the same amount is spread over a longer period, the consequences are not so severe. How many time between the twelve drinks is needed to not reach the risk zone? One suitable answer is provided by COGORZA. With a drinking time of an hour and a waiting time of an other hour, a male drinker of 92 kg weight with can drink twelve cups of wine in a full day without reaching drunkenness state(Figure 5).

**Figure 5.**
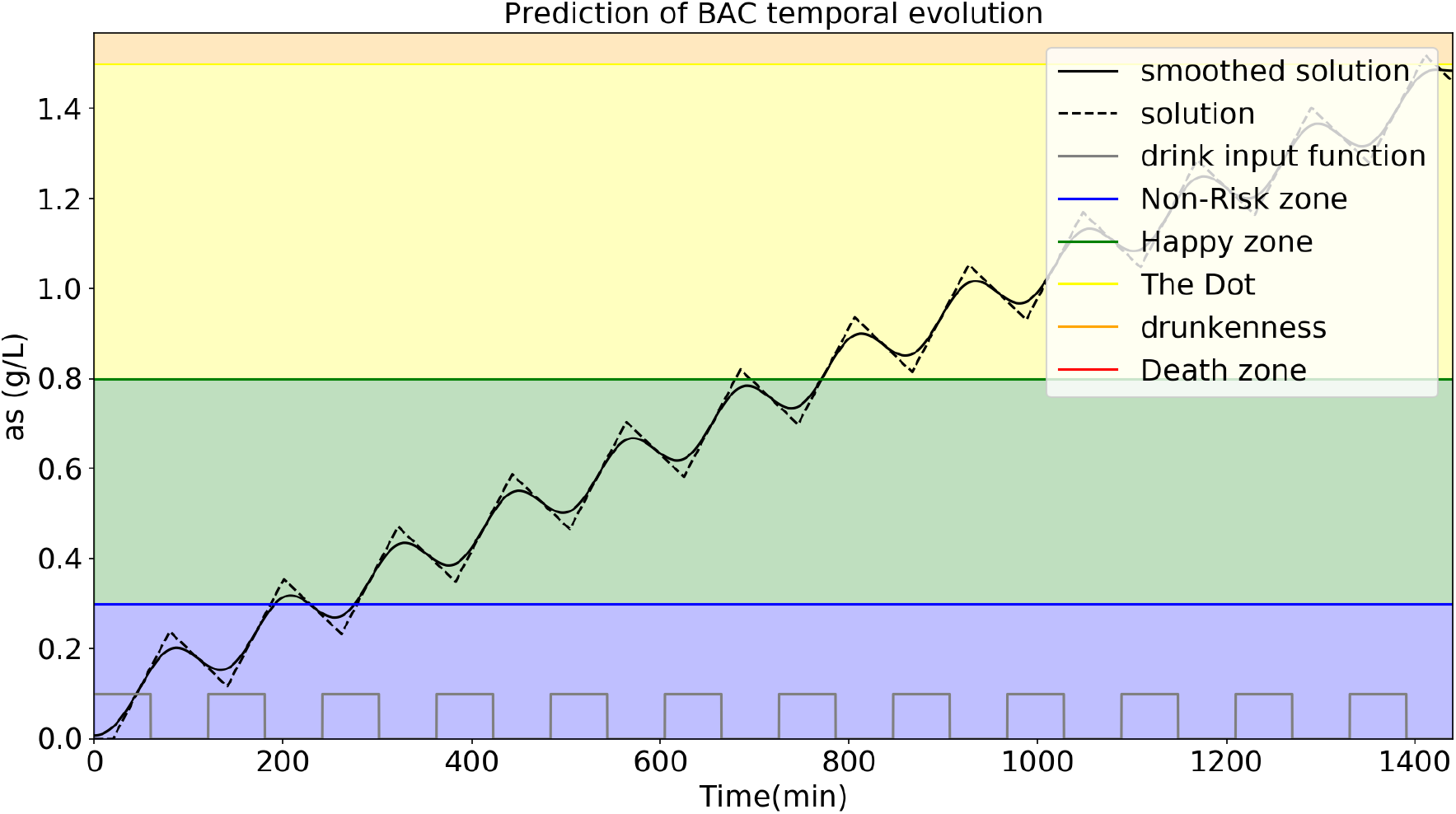
Predicted temporal BAC evolution for third study case. This case represents a drinker that drinks one cup of wine every two hours with an interruption time in between of one hour (black dotted line). The solid black line represents the smoothed solution. The step functions in gray in the lower part represents the input drinks made. Colored areas in the background represent areas of risk due to BAC levels.

## 5 Limitations

This model is exposed to the general public and especially to the specialist for its initial assessment or experimental contrast. It must be considered that this model is based on some magnitudes that for simplicity are set as constants or approximated with a simple expression. The proposed values can be modified in order to adapt the code to other situations or to make a better approximation to the real solution. For instance, the total amount of blood could be modelled using gender, body mass and height as did in [4].

One of the limitations of this model is the assumption that the drinker is continuously consuming a particular drink from the moment they start. This limitation can be partially overcome if instead of considering a step function for each drink, is performed by considering each sip. However, this approach requires a good characterization of the time sequence of sips, as well as the amount ingested in each of them. Additional adjustments can be made taking into account the size of the glass, the presence and amount of ice or the proportion of alcohol in case it is a mixture (for example, a free tank). In the case of application to other scenarios, such as intake of medicines, the input waveform would be computed from the number of pills consumed and the amount of mass of interest present in each pill.

The ramp-type solution obtained is due to input nature defined as step-wise functions. The input functions can be defined in other ways, as for instance by Gaussian kernels, which would give smoothed results. If so, a constraining of the total mass amount of a given drink must be conserved and distributed between initial and final times. The smoothing of the solution is a way to partially solve this inconvenience, but it becomes quite inaccurate near to time boundaries. For instance, in the analyzed cases in this work, alcohol level immediately pass to bloodstream after start drink. From what is known, alcohol enters in blood after 10 minutes since drink starts.

A remarkable limitation of the work is that alcohol elimination by liver is take as the only way, while is known that 90% is removed by liver and the rest by lungs and kidneys [9]. That limitation can be sorted out by applying a factor 1.1 to elimination rate in equation 5.

## 6 Conclusions

The main motivation I had to start developing this work was the aim to understand the phenomenology underlying the time evolution of the concentration of ingested substances in blood. It is common that even experienced drinkers do not have effective tools beyond their own experience to determine the potential intoxication that would produce a set of drinks in a given sequence of time [10, 11]. For this reason, a prior planning and scenario analysis are useful to anticipate and be aware of the expected outcome of potential alcohol abuse.

The purpose of this model is to shed some light on the prediction of the blood alcohol curve that can be given from a series of alcoholic drinks especially considering timing input configuration. The proposed model presented in this work is able to predict a BAC evolution which is consistent with what is known about the effects of substances in the body. The model presented predicts BAC peak and the time required for complete body cleaning.

One of the relevant points of the presented work is it’s pedagogical nature. The proposed model has a potential application for the analysis of substance curves in blood based on conservation principles and knowledge about the absorption and elimination rates. It is hoped that the reader will use the model to better understand BAC evolution for a given series of drinks.

## 7 Source code

In the following sections is exposed the source code developed to implement the theoretical model proposed. The code has been written in *python*, where a male subject of M=92 kg of body mass ingests twelve alcoholic drinks (i.e. twelve bottles of beer) in 12 h. Total simulation time is 24 h.

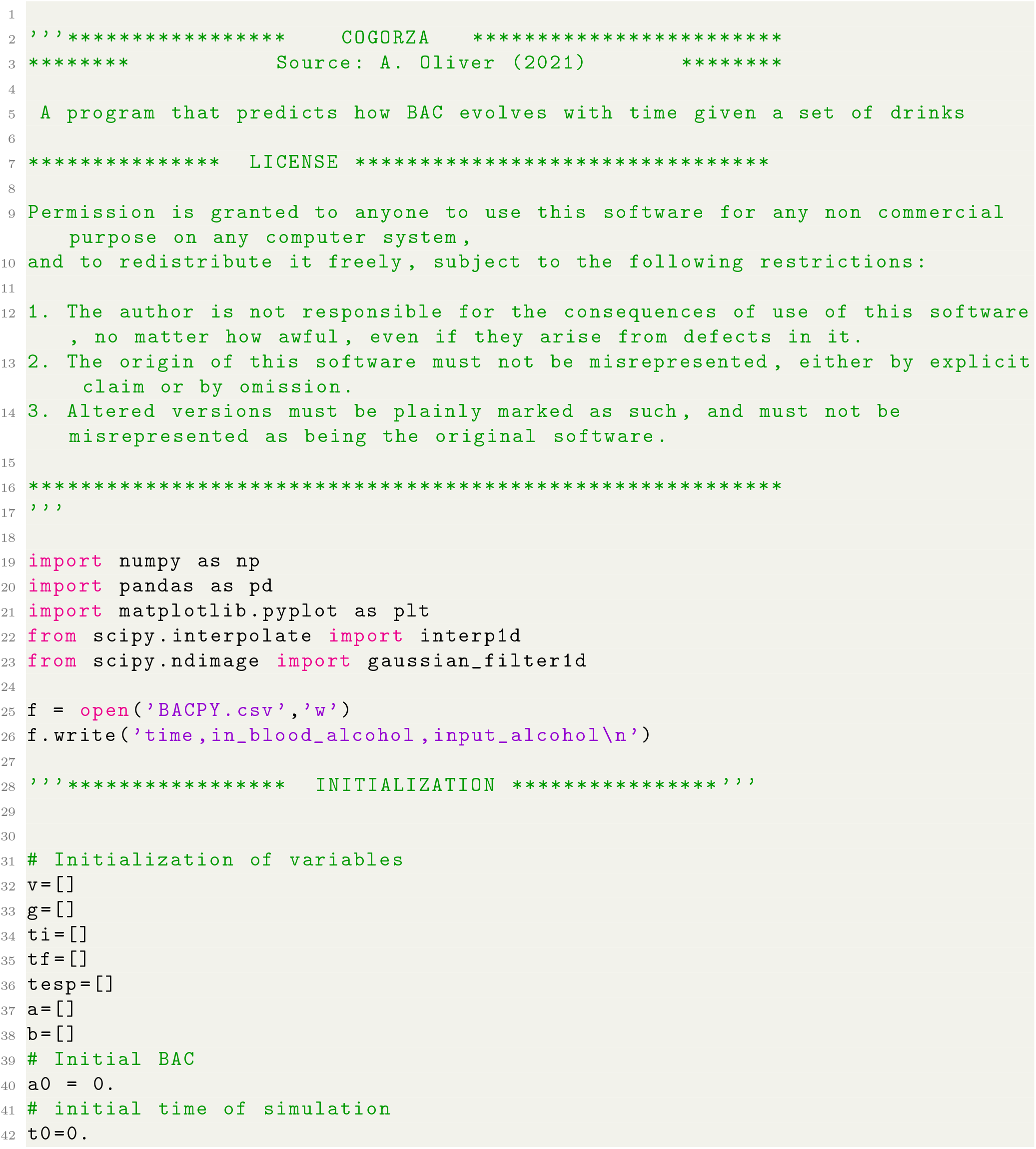

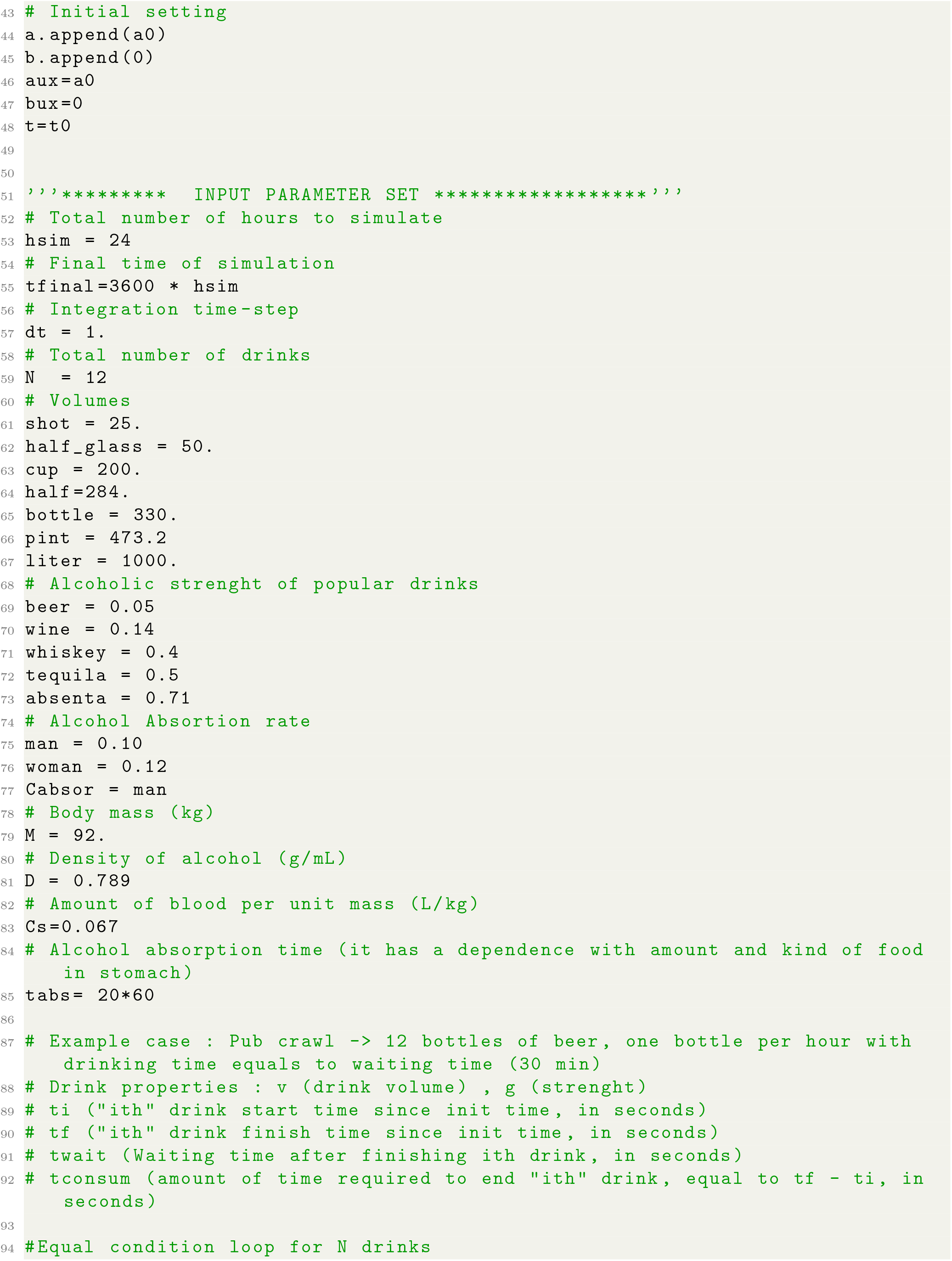

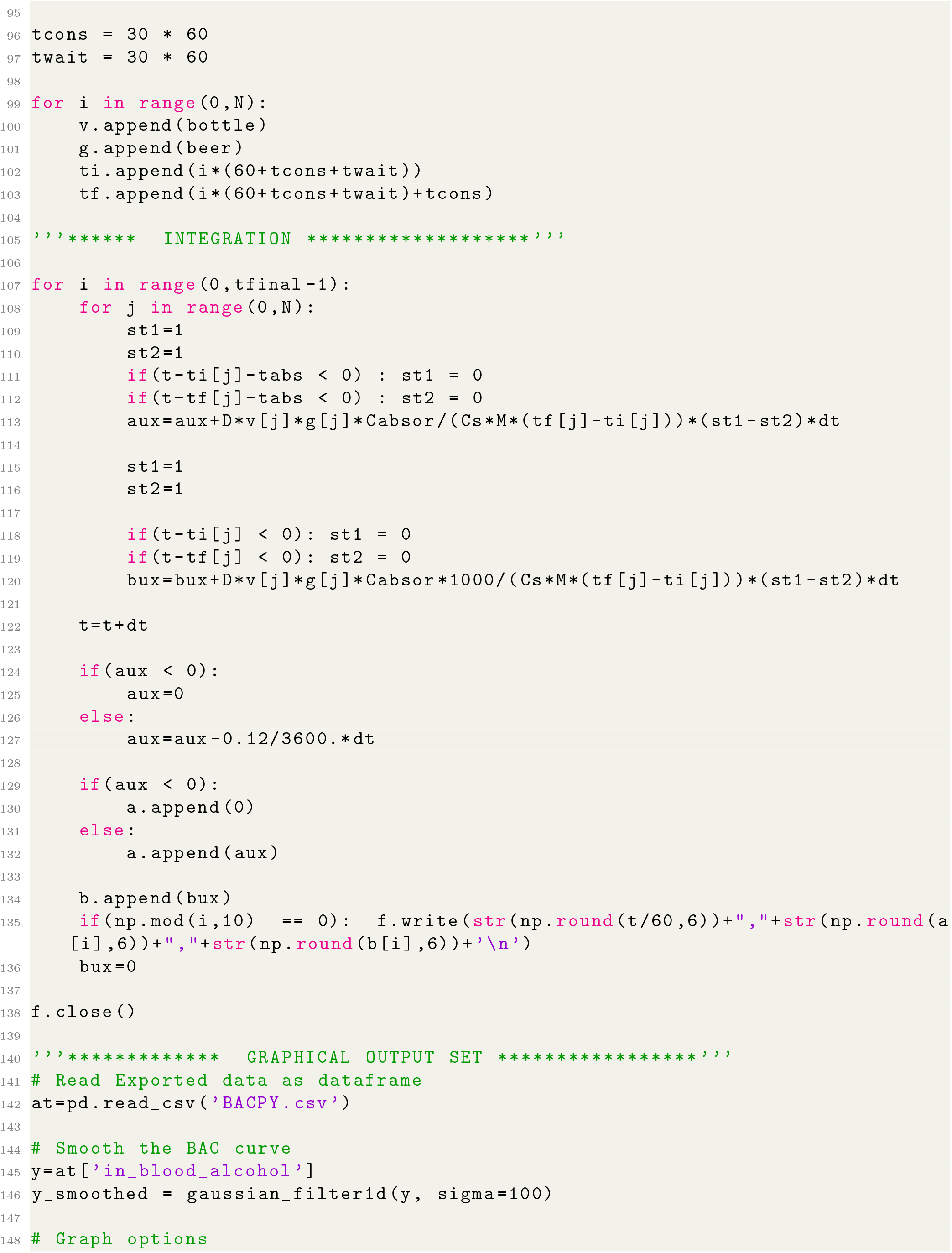

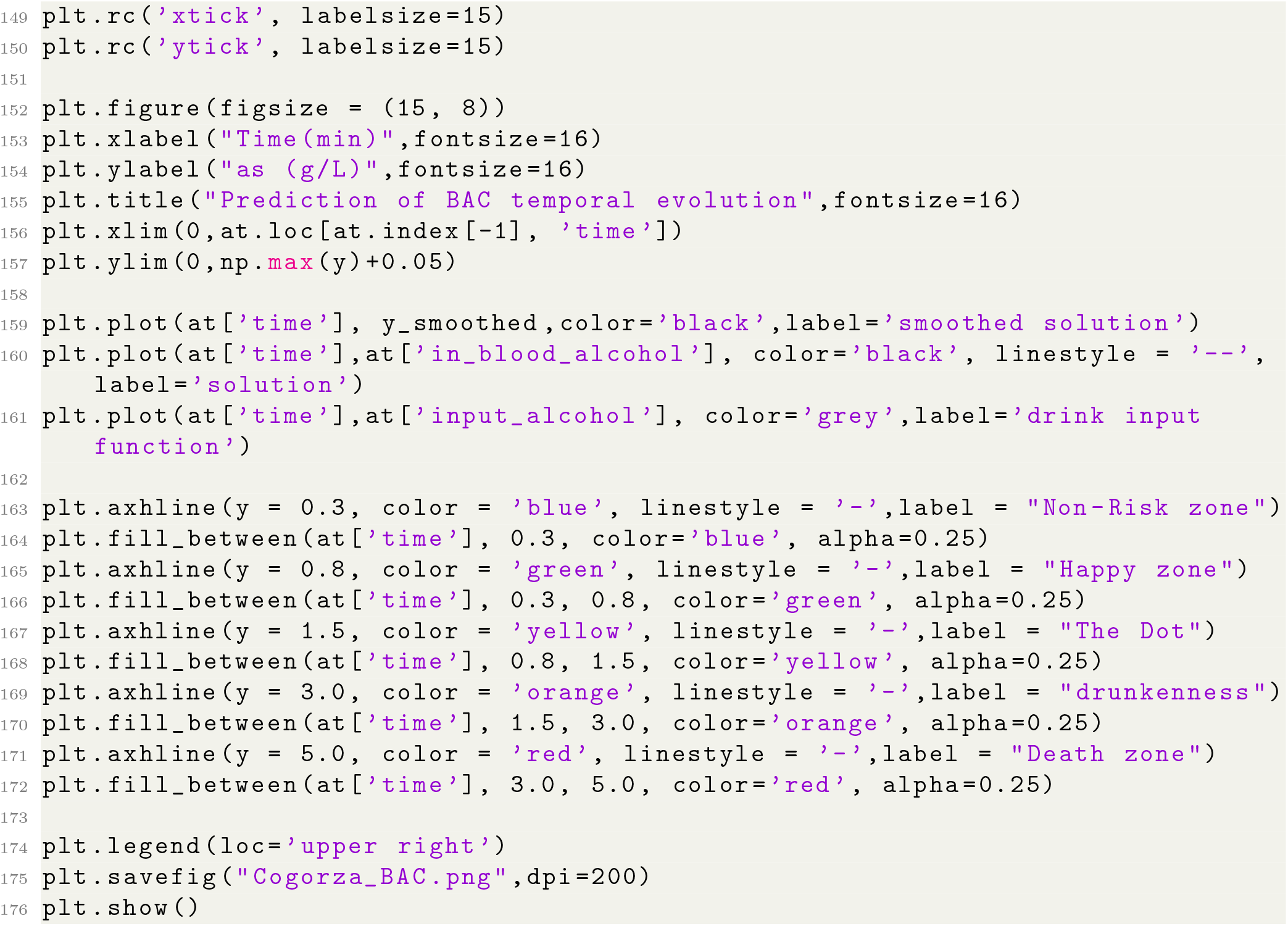

## Footnotes

2 https://www.slideshare.net/6yinyan9/absorcin-de-alcohol

3 https://estilosdevidasaludable.sanidad.gob.es/consumo/falsosMitos/trucos/home.htm

4 http://www.ieslaasuncion.org/fisicaquimica/tasa.htm

5 http://vitacoratxc.blogspot.com/2008/03/absorcin-y-eliminacin-del-alcohol-los.html

